# A single cell transcriptome atlas reveals the heterogeneity of the healthy human cornea and identifies novel markers of the corneal limbus and stroma

**DOI:** 10.1101/2021.07.07.451489

**Authors:** Pere Català, Nathalie Groen, Jasmin A. Dehnen, Eduardo Soares, Arianne J.H. van Velthoven, Rudy M.M.A. Nuijts, Mor M. Dickman, Vanessa L.S. LaPointe

## Abstract

The cornea is the clear window that lets light into the eye. It is composed of five layers: epithelium, Bowman’s layer, stroma, Descemet’s membrane and endothelium. The maintenance of its structure and transparency are determined by the functions of the different cell types populating each layer. Attempts to regenerate corneal tissue and understand disease conditions requires knowledge of how cell profiles vary across this heterogeneous tissue. We performed a single cell transcriptomic profiling of 19,472 cells isolated from eight healthy donor corneas. Our analysis delineates the heterogeneity of the corneal layers by identifying cell populations and revealing cell states that contribute in preserving corneal homeostasis. We identified that the expression of *CAV1*, *CXCL14*, *HOMER3* and *CPVL* were exclusive to the corneal epithelial limbal stem cell niche, *CKS2*, *STMN1* and *UBE2C* were exclusively expressed in highly proliferative transit amplifying cells, and *NNMT* was exclusively expressed by stromal keratocytes. Overall, this research provides a basis to improve current primary cell expansion protocols, for future profiling of corneal disease states, to help guide pluripotent stem cells into different corneal lineages, and to understand how engineered substrates affect corneal cells to improve regenerative therapies.

## INTRODUCTION

The healthy cornea is a transparent and avascular tissue that allows light to enter the eye and accounts for most of its refractive power. The cornea is composed of five layers: its outer surface is a stratified sheet of corneal epithelial cells that reside on the Bowman’s layer, a collagen-based acellular membrane synthesized by the stromal keratocytes. The keratocytes populate the corneal stroma, the middle layer of the cornea that is composed of structured collagen fibers and other extracellular matrix proteins. The corneal endothelium forms a thin monolayer of tightly packed hexagonal cells that line in the innermost surface of the cornea and reside in close contact to the stroma on the Descemet’s membrane.

Corneal structure and transparency are governed by the functions of the cell types populating each layer. Epithelial cells act as a biological barrier to block the passage of foreign material and provide a smooth surface that absorbs nutrients. Keratocytes maintain extracellular matrix homeostasis responsible for the cornea’s biomechanical and optical properties, and endothelial cells serve as active pumps transporting ions, metabolites and fluid to maintain corneal hydration and transparency.

Selective replacement of dysfunctional single corneal layers with that of a donor,^1^ the autologous transplantation of primary cultured corneal epithelial limbal stem cells,^2^ and the treatment of endothelial dysfunctions with primary cultured allogeneic corneal endothelial cells^3^ are already a therapeutic reality. Gaining deeper understanding of corneal cell profiles and their transcriptomic signatures can be highly relevant for the improvement of such therapies.

In order to further understand the cellular complexity of this heterogeneous tissue, we provide a high-quality single-cell ribonucleic acid sequencing (scRNAseq) dataset from 19,472 corneal cells isolated from four female and four male donors. With it, we depict the diverse cell populations and provide a comprehensive cell atlas of the healthy human cornea. The transcriptomic cell census identified subpopulations with different roles in the maintenance of corneal homeostasis and provides a baseline to improve primary cell expansion protocols, for future profiling of corneal disease states, to help guide pluripotent stem cells into different corneal lineages, and to understand how engineered substrates affect corneal cells to improve regenerative therapies. Furthermore, this dataset identified markers exclusively expressed in cells comprising the limbal epithelial stem cell niche, highly proliferating epithelial cells, and stromal keratocytes which could be used as reference to improve current corneal cell replacement therapies.

## RESULTS

### Five major cell clusters were identified in the corneal tissue

Six donor corneas and the limbi of two donor corneas (Table 1) were manually dissected and dissociated into single cells for single-cell RNA sequencing. After filtering low quality cells, the transcriptome profiles of 19,472 cells were further analyzed (Figure 1a).

**Figure 1.**
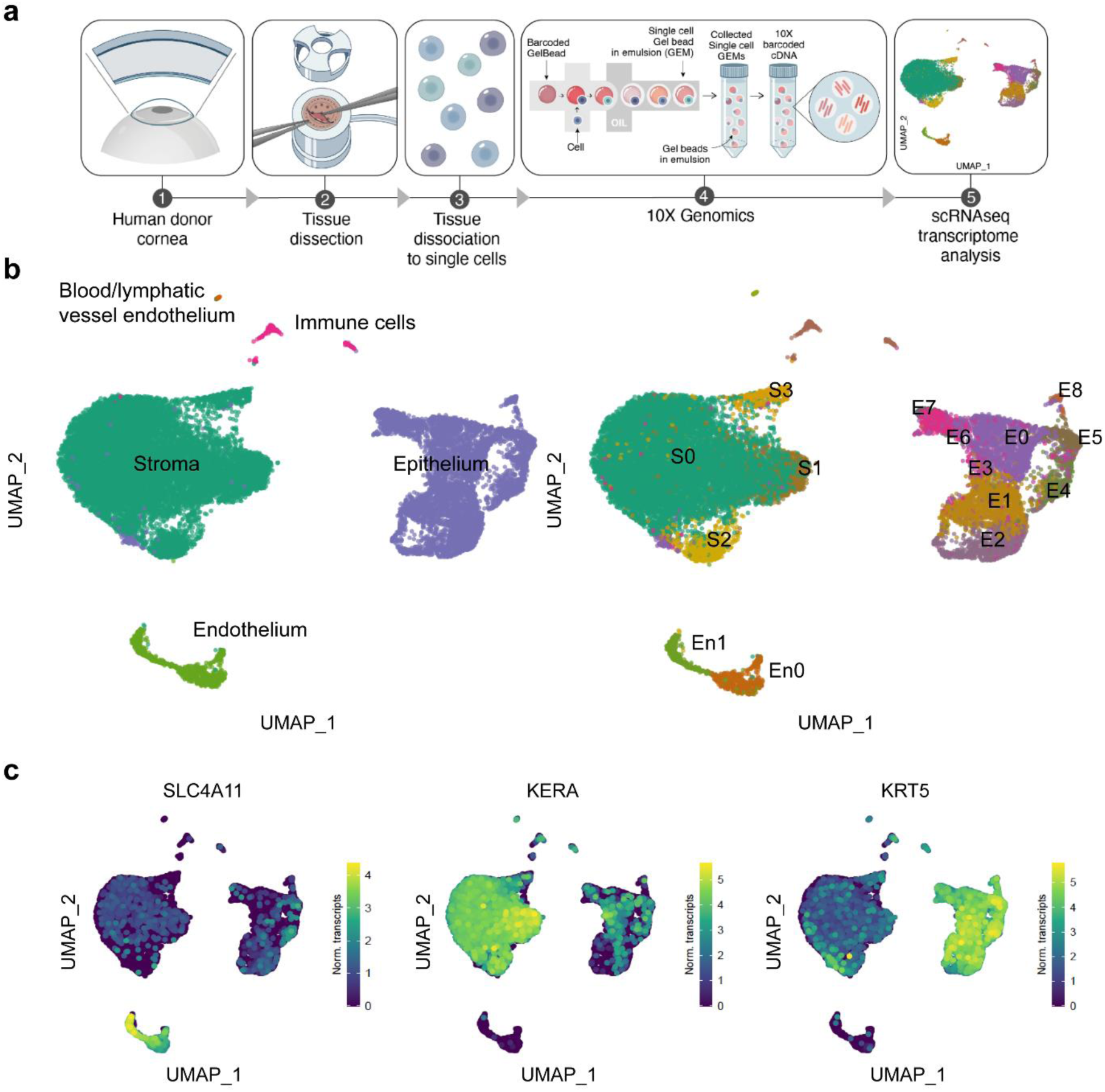
Cell clusters from all three corneal layers and two additional clusters of immune and blood/lymphatic endothelial cells were identified with scRNAseq analysis. Experimental overview for scRNAseq (a). UMAP of the 19,472 sequenced cells with the identified five major cell clusters (b, left). Cell clusters from the three corneal cell layers were further subclustered in different cell subpopulations (b, right). Single cell expression level UMAP of specific corneal layer markers supports cell cluster identification (c): *SLC4A11* is corneal endothelial specific, *KERA* is corneal stromal specific, and *KRT5* is corneal epithelial specific.

**Table 1.** Donor cornea information

| Cornea | Gender | Age (years) | Preservation | Eye bank | Experiment |
| --- | --- | --- | --- | --- | --- |
| 01 | Female | 62 | Optisol (4°C for 5 days) | Saving Sight | Full cornea scRNA-seq |
| 02 | Male | 22 | Optisol (4°C for 6 days) | Saving Sight | Full cornea scRNA-seq |
| 03 | Male | 75 | Optisol (4°C for 10 days) | Saving Sight | Full cornea scRNA-seq |
| 04 | Female | 48 | Optisol (4°C for 8 days) | Saving Sight | Full cornea scRNA-seq |
| 05 | Male | 75 | Organ culture media (31°C for 21 days) | ETB-BISLIFE | Full cornea scRNA-seq |
| 06 | Female | 75 | Organ culture media (31°C for 15 days) | ETB-BISLIFE | Full cornea scRNA-seq |
| 07 | Female | 71 | Organ culture media (31°C for 18 days) | ETB-BISLIFE | Limbus scRNA-seq |
| 08 | Male | 79 | Organ culture media (31°C for 14 days) | ETB-BISLIFE | Limbus scRNA-seq |
| 09 | Female | 78 | Organ culture media (31°C for 7 days) | ETB-BISLIFE | Immunofluorescence |
| 10 | Male | 71 | Organ culture media (31°C for 8 days) | ETB-BISLIFE | Immunofluorescence |

Data of the 19,472 sequenced cells were embedded in a uniform manifold approximation and projection (UMAP) and unbiased low resolution clustering revealed five major cell clusters (Figure 1b, left). The expression of specific corneal layer marker (keratin 5 (*KRT5)* for epithelium^4^, keratocan (*KERA)* for stromal cells^5^, and transporter-like protein 11 (*SLC4A11)* for endothelium^6^) identified the three main layers within the five identified clusters (Figure 1c). Differential gene expression profiling of the clusters further confirmed the identification of a corneal epithelial cell cluster comprising 5,964, a corneal stromal cell cluster comprising 12,344 cells, a corneal endothelial cell cluster comprising 842 cell, and non-corneal clusters of blood/lymphatic vessel endothelial cells comprising 36 cells, and immune cells comprising 216 cells (Figure 2). Cell clusters from the three corneal cell layers were further classified separately at higher resolution in different corneal layer–specific subclusters (Figure 1b, right).

**Figure 2.**
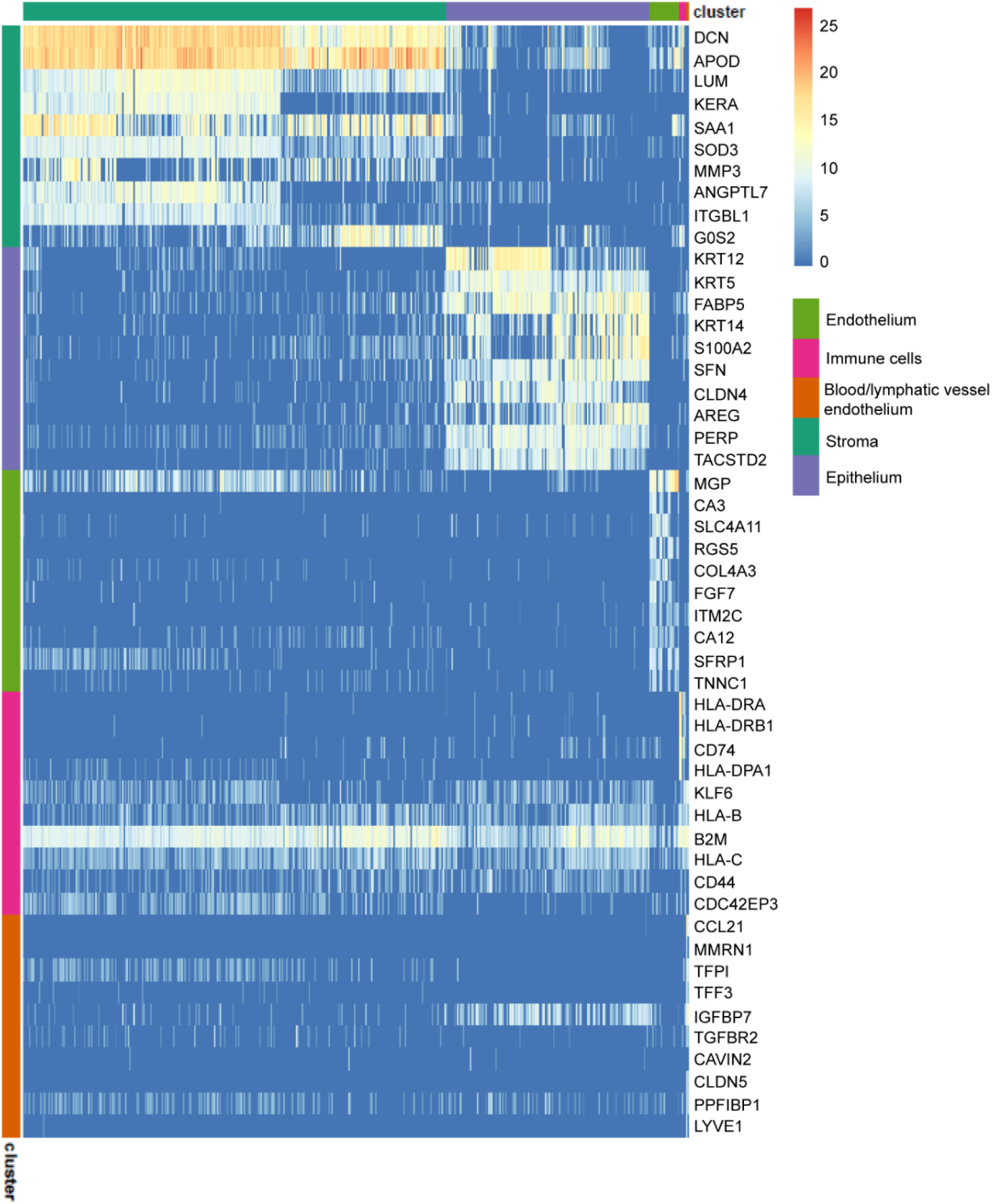
The heatmap of the top 10 differentially expressed genes for each cluster showed distinct transcriptomic profiles for the five major cell clusters and allowed cell cluster identification.

### Nine cell clusters were identified within the corneal epithelium

A major cluster of 5,964 corneal epithelial cells was identified and further analysis revealed nine subclusters (E0-8, Figure 3a). Differential gene expression profiling was used for further identification of the subclusters. Cluster E3 presented a high expression of stromal cell markers *KERA*, lumican (*LUM*), and aldehyde dehydrogenase 3 member A1 (*ALDH3A1*),^5, 7, 8^ as well as a reduced expression of corneal epithelial marker keratin 12 (*KRT12*)^9, 10^ compared to the other epithelial subclusters (Figure 3b and Supplementary Figure S1). Putative doublet analysis of cluster E3 suggested epithelial–stromal keratocyte cell doublets (Supplementary Figure S2). Cluster E7 was identified as conjunctival epithelial cells based on high expression of conjunctival markers such as keratin 13 (*KRT13*), keratin 15 (KRT15), and keratin 19 (*KRT19*),^11, 12^ and the low expression of the corneal epithelial marker *KRT12* (Figure 3b and Supplementary Figure S1).

**Figure 3.**
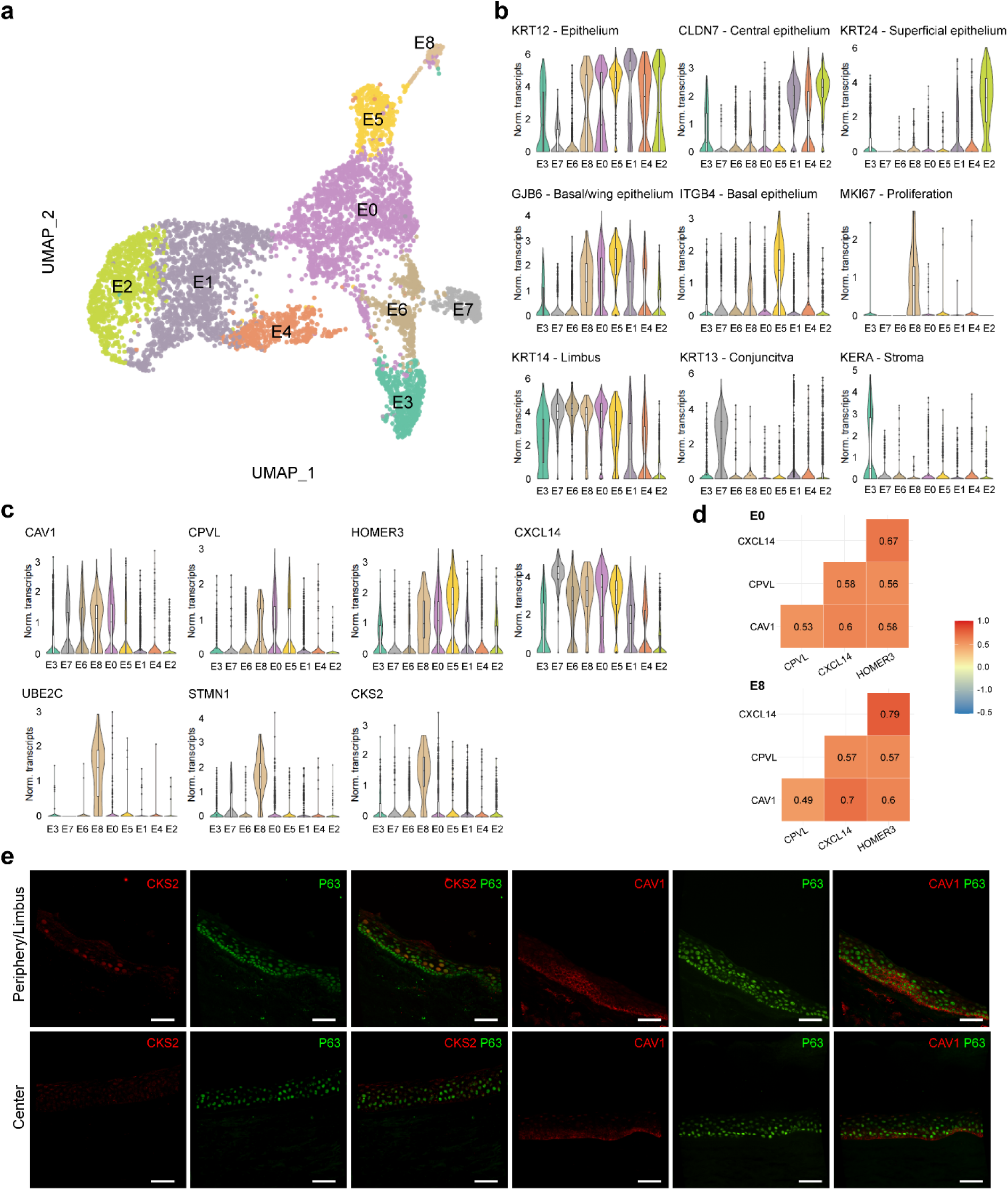
A major cluster of 5,964 corneal epithelial cells was identified with scRNAseq analysis. The UMAP revealed nine different cell subclusters: a cluster of basal limbal epithelial cells (E0), a cluster of migratory epithelial cells (E1), a cluster of wing/superficial epithelial cells (E2), a cluster of stromal-epithelial doublets (E3), a cluster of central epithelial cells (E4), a cluster of basal central epithelial cells originating from the limbus (E5), a cluster of wing epithelial cells in the limbus (E6), a cluster of conjunctival cells (E7), and a cluster of highly proliferative transit-amplifying cell (E8) (a). Violin plots show marker genes for corneal epithelium (KRT12), central (*CLDN7*), superficial (*KRT24*), suprabasal (*GJB6*), basal (*ITGB4*), and limbal (*KRT14*) corneal epithelium, as well as proliferation (*MKI67*), conjunctival (*KRT13*) and stromal (*KERA*) markers, for the identification of cell subclusters (b). Differential gene expression identified novel epithelial limbal niche–specific markers and proliferative epithelial cell–specific markers, and are visualized as violin plots (c). Correlation analysis confirmed the co-expression of limbal niche–specific markers in peripheral/limbal epithelial cell populations (d). Immunofluorescence of *CKS2* and caveolin-1 (*CAV1*) (red) on human corneal tissue cryosections confirmed differential protein expression in the limbus/periphery (e, top row), and absence (*CKS2*) or minimal expression (caveolin-1) in the central cornea (e, bottom row). P63 (ΔNp63 or p63α) was used as corneal epithelial limbal cell marker (green). Basal corneal epithelial cells retained minimal expression of both ΔNp63 and caveolin-1, suggesting their limbal origin. Scale bar is 50 μm.

Cells forming clusters E6, E8 and E8 showed increased expression of corneal limbal markers keratin 14 (*KRT14*)^13, 14^, *KRT15*,^15, 16^ and S100 calcium binding protein A2 (*S100A2)*^17^ compared to the other identified clusters suggesting their location in the corneal limbus (Figure 3b and Supplementary Figure S1). Interestingly, cluster E6 showed an increased expression of the superficial epithelium limbus marker^17^ S100 calcium binding protein A8 (*S100A8*) compared with clusters E0 and E8 and a reduced expression of the basal corneal epithelial cell markers connexin 26 (*GJB2*), connexin 30 (*GJB6*), and integrin β4 (*ITGB4*),^18–20^ (Figure 3b and Supplementary Figure S1) leading to its identification as a population of wing/superficial epithelial cells in the limbus or peripheral cornea. Cluster E0 presented an increased expression of *GJB6*, and *GJB2*, which are predominantly found in basal corneal epithelium^18^ suggesting these cells formed a basal corneal epithelial cell population in the limbal stem cell niche or peripheral cornea (Figure 3b and Supplementary Figure S1). Finally, the high expression of mitogenic factors Ki-67 (*MKI67*)^21^, survivin (*BIRC5*)^22^ and H2A histone family member X (*H2AFX*)^23^ in cluster E8, as well as the differential expression of transit-amplifying cell marker CD109^24^, suggested that cluster E8 was formed by highly proliferative transit-amplifying cells in the limbal stem cell niche or peripheral cornea (Figure 3b and Supplementary Figure S1). No quiescent limbal epithelial stem cells expressing *ABCB5*, *ABCG2* and *CD200*^24–27^ were identified in this dataset (Supplementary Figure S1).

Cluster E5 presented high expression of *GJB6* and *ITGB4* associated with basal epithelium^18^ and *KRT12,* and *KRT3* associated with terminally differentiated corneal epithelium^27–29^, along with a reduced expression of *KRT14* compared to clusters E6, E8, and E0, and no expression of *KRT15,* suggesting these cells were corneal epithelial basal cells originating from the limbus (Figure 3b and Supplementary Figure S1). Cluster E1 was identified as post-mitotic and terminally differentiated migratory epithelial cells based on the expression of genes associated with cell migration such as *RHOV*^30^ and tight junction formation and obliteration *CLDN7*^31^ together with a high expression of corneal epithelial cell markers *KRT12*, *KRT3,* and *KRT5*. This cluster retained a low expression level of *KRT14*, implying their limbal origin (Figure 3b and Supplementary Figure S1).

Cluster E4 was composed of cells with a high expression of *KRT12, KRT3*, and *KLF5* suggesting this cluster was terminally differentiated cells from the central corneal epithelium (Figure 3b and Supplementary Figure S1). The cells forming cluster E2 presented a high expression of *KRT12*, *KRT24*, and *CXCL17* associated with wing/superficial central epithelium (Figure 3b and Supplementary Figure S1).

### ScRNAseq reveals novel specific markers for the corneal limbal stem cell niche and transit-amplifying cells

Differential gene expression analysis of the basal corneal limbal epithelial cells (cluster E0), and the transit-amplifying cells (cluster E8), showed that genes encoding for caveolin-1 (*CAV1*), probable serine carboxypeptidase (*CPVL*), homer scaffolding protein 3 (*HOMER3*), and C-X-C motif chemokine 14 (*CXCL14*) were highly expressed in both clusters (Figure 3c), suggesting they could be markers of the human limbal stem cell niche. Moreover, the expression of the markers highly correlated in clusters E0 and E8 (Figure 3d), again suggesting their identity of corneal epithelial cells located in the limbal stem cell niche. Interestingly, a cluster of basal corneal epithelial cells (cluster E5) retained some expression of the identified markers, in line with the limbal origin annotated for this cluster. To confirm this, caveolin-1 (*CAV1*) expression in the corneal epithelial limbus was assessed by immunofluorescence (Figure 3e) and further validated on human primary cultured corneal limbal epithelial cells, where it was expressed in ΔNp63-positive limbal epithelial stem cells (Supplementary Figure S3). Differential gene expression analysis on the transit-amplifying cells (cluster E8) revealed that the expression of cyclin-dependent kinase 2 (*CKS2*), stathmin (*STMN1*), and ubiquitin conjugating enzyme E2 C (*UBE2C*) was exclusive to this cluster. Cyclin-dependent kinase 2 (*CKS2*) expression in the corneal epithelial limbus was confirmed by immunofluorescence (Figure 3e) and further validated on human primary cultured corneal limbal epithelial cells, where it was expressed in in p63α-positive limbal epithelial stem cells (Supplementary Figure S4).

### Four cell clusters were identified within the corneal stroma

A major cluster of 12,344 corneal stromal cells was identified, and further analysis revealed four different subclusters of corneal stromal cells (S0-S3, Figure 4a). High expression of stromal keratocyte markers LUM, *KERA*, or *ALDH3A1*^5, 7, 8^ suggested that clusters S0, S1 and S2 were stromal keratocytes (Figure 4b). Cluster S2 showed increased expression of extracellular matrix proteins such as *COL12A1*, *COL6A2*, *COL6A1*, and *LAMB2* suggesting these were activated keratocytes that played a crucial role in maintaining the corneal stromal extracellular matrix (Figure 4b and Supplementary Figure S5). Cluster S3 showed a decreased expression of keratocyte markers *LUM, KERA*, and *ALDH3A1* and increased expression of the fibroblastic marker *CD44*^32–34^ compared to clusters S0, S1 and S2 (Figure 4b and Supplementary Figure S5). The myofibroblast-specific marker, α-smooth muscle actin (*ACTA2*),^35, 36^ was not detected in cluster S3 (Supplementary Figure S5), suggesting it is composed of keratocytes that are in transition to stromal myofibroblasts.

**Figure 4.**
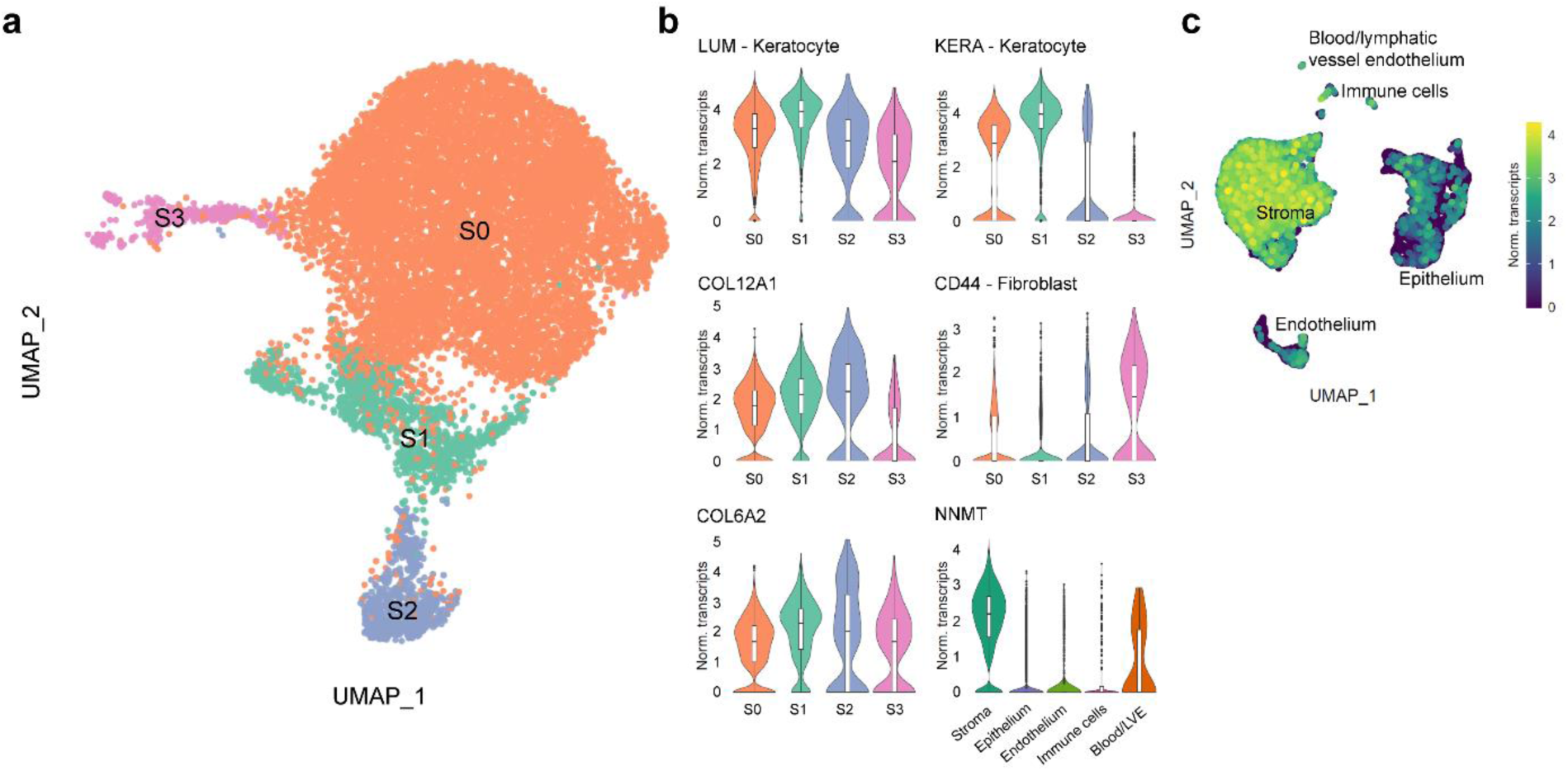
A major cluster of 12,344 corneal stromal cells was identified with scRNAseq analysis. The UMAP revealed four different cell subclusters, three keratocyte clusters (S0-S2), with one having a high extracellular matrix protein secretion profile (S2) and a cluster of keratocytes transitioning to stromal myofibroblasts (S3) (a). Violin plots show marker genes for stromal keratocytes (*KERA* and *LUM*), stromal fibroblasts (*CD44*), collagen (*COL6A2* and *COL12A1*) secretion for the identification of cell subclusters, and *NNMT* expression exclusive to the stroma within the cornea (b). Single cell expression level UMAP of NNMT further confirms differential expression of *NNMT* in the stromal cluster (c).

Finally, differential gene expression analysis of the major stromal cluster compared to the clusters associated with the corneal epithelial and endothelial layers identified that the expression of nicotinamide N-methyltransferase (*NNMT*) was exclusive to the corneal stromal cluster, suggesting that this gene could be used as a novel corneal stromal marker (Figure 4b and c).

### Two cell clusters were identified within the corneal endothelium

A major cluster of 842 corneal endothelial cells was identified and further analysis revealed two different subclusters of corneal endothelial cells (En0-1, Figure 5a). Both clusters showed expression of corneal endothelial cell markers CD166 (*ALCAM*) and sPrdx1 (*PRDX1*)^37, 38^ and functional markers Na^+^/K^+^ ATPase (*ATP1A1*)^6^ and sodium bicarbonate transporter-like protein 11 (*SLC4A11*)^6^ confirming the corneal endothelial phenotype (Figure 5b and Supplementary Figure S6). Differential expression analysis revealed that cluster En0 possessed a lower expression of tight junction protein zona occludens-1 (*TJP1*)^6^ and focal adhesion regulator microtubule-actin cross-linking factor-1 (*MACF1*)^39^ compared to cluster En1 (Figure 5b), suggesting cells in cluster En0 could preferentially migrate upon corneal endothelial damage to contribute to tissue repair. Furthermore, cluster En1 possessed a higher expression of *COL4A3* (Figure 5b) suggesting these cells could play an important role on Descemet’s membrane homeostasis. Interestingly, both clusters retained expression of *PITX2*, a periocular mesenchyme marker associated with progenitor endothelial cells.^40, 41^ No endothelial fibroblasts were identified, evidenced by the absence of *ACTA2* and *CD44* (Supplementary Figure S6).

**Figure 5.**
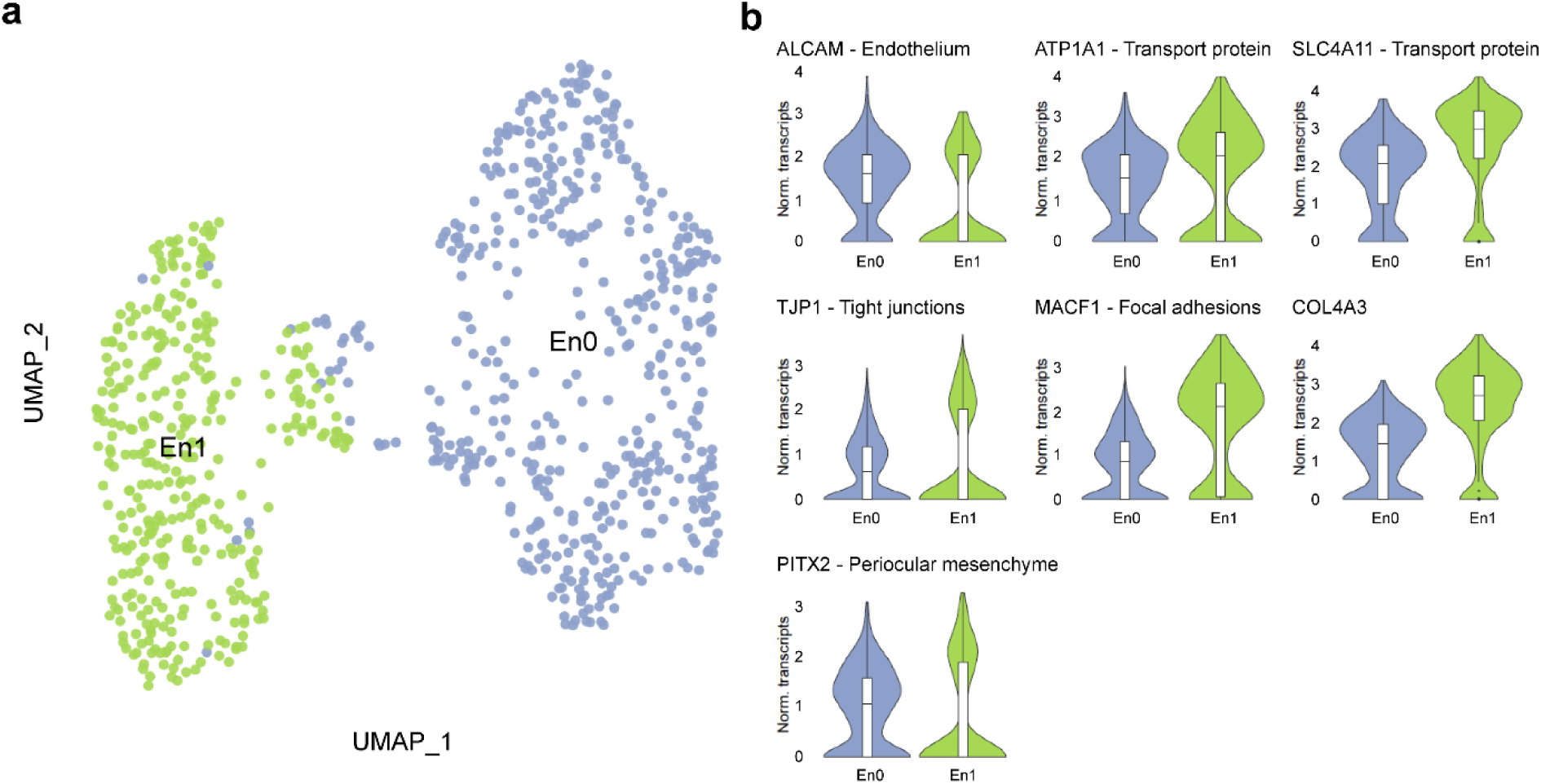
A major cluster of 842 corneal endothelial cells was identified with scRNAseq analysis. The UMAP revealed two different endothelial cell subclusters, a cell cluster with high collagen synthesis (En2) and another cell cluster with low tight junction and focal adhesion protein expression (En1) (a). Violin plots show marker genes for corneal endothelium (*ALCAM*), ion and bicarbonate transporters (*ATP1A1* and *SLC4A11* respectively), tight junction proteins (*TJP1*), focal adhesion protein regulator (*MACF1*), collagen secretion (*COL4A3*) and periocular mesenchyme (*PITX2*) (b).

## DISCUSSION

Gaining transcriptomic information at the single cell level of human corneal cells enables a greater understanding of this heterogeneous tissue. In this study, we have performed a scRNAseq analysis of the healthy cornea to create a comprehensive cell atlas of the human cornea (Figure 6). Moreover, scRNAseq analysis enabled the identification of novel markers of the limbal epithelial stem cell niche, transit-amplifying cells, and stromal keratocytes. The data generated can serve as a reference cell atlas with a major impact in the further improvement and development of cell replacement therapies or regenerative medicine approaches for treating corneal blindness. This research further complements scRNAseq analysis of the developing human cornea^42^ and of the human corneal limbus^43^.

**Figure 6.**
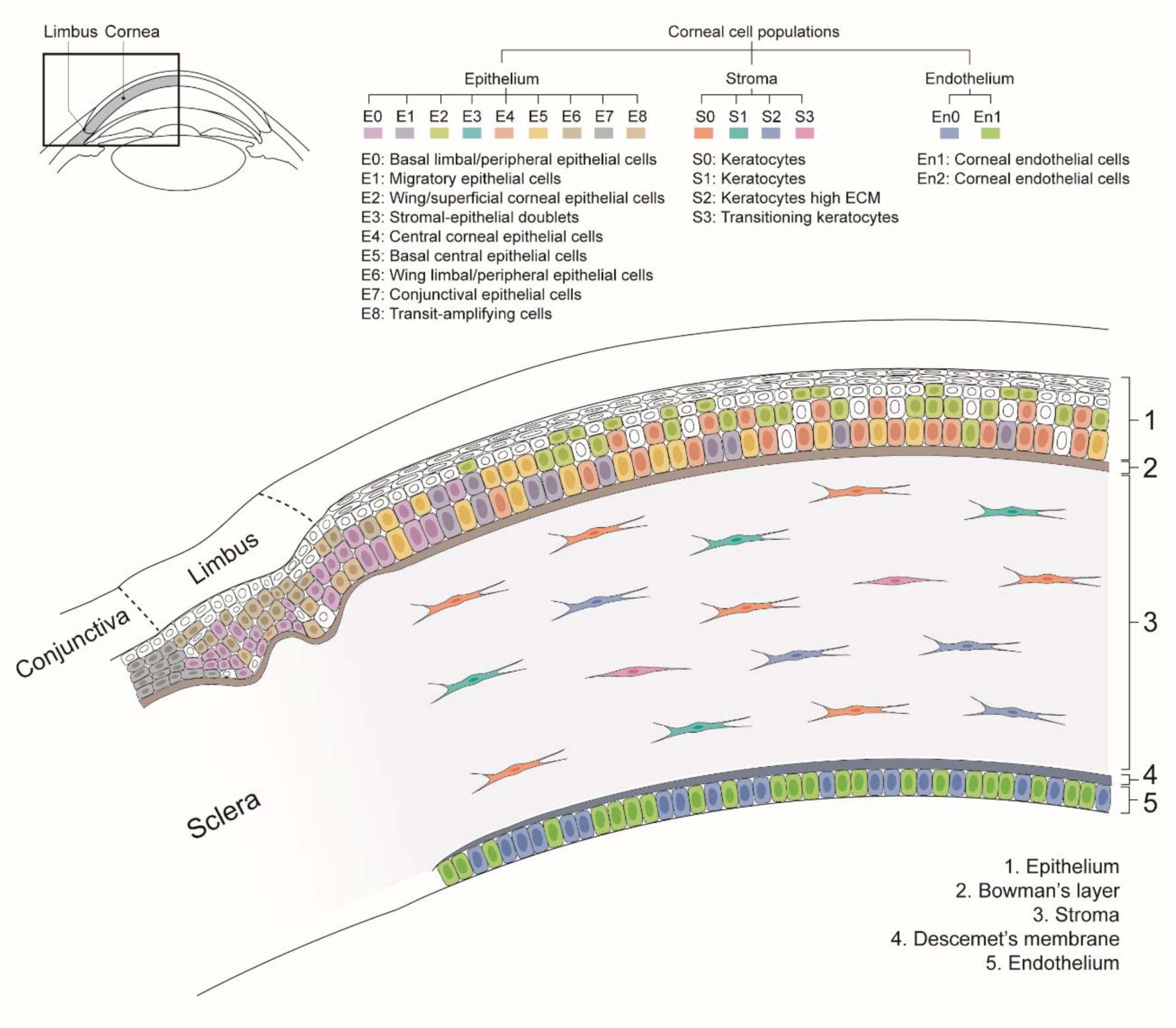
scRNAseq data analysis identified nine epithelial, four stromal and two endothelial corneal cell clusters. This figure presents a summary of the cell type and location of each cluster in the healthy human cornea, generating a comprehensive cell census of the healthy human cornea. In the epithelium: a cluster of basal limbal epithelial cells (E0), a cluster of migratory epithelial cells (E1), a cluster of wing/superficial epithelial cells (E2), a cluster of stromal-epithelial doublets (E3), a cluster of central epithelial cells (E4), a cluster of basal central epithelial cells originating from the limbus (E5), a cluster of wing epithelial cells in the limbus (E6), a cluster of conjunctival cells (E7), and a cluster of highly proliferative transit-amplifying cell (E8). In the stroma: three keratocyte clusters (S0-S2), with one having a high extracellular matrix protein secretion profile (S2) and a cluster of keratocytes transitioning to stromal myofibroblasts (S3). In the endothelium: a cell cluster with high collagen synthesis (En2), and a cell cluster with low tight junction and focal adhesion protein expression (En1).

The corneal epithelium appeared to be the most heterogeneous corneal layer with 9 identified cell clusters. The transcriptomic signature of epithelial cells from the basal, wing and superficial layers of both central and limbus/peripheral cornea were identified. A population of conjunctival epithelial cells was detected (cluster E7; Figure 3), likely representing contamination from the dissection process. Moreover, a cluster of highly proliferative transit-amplifying cells (cluster E8) was detected. No quiescent limbal epithelial stem cells expressing *ABCB5, TCF4, CD200* and *ABCG2* were identified (Supplementary Figure S1). This is in line with the study of Li et al., where out of 16,360 cells specifically isolated from the adult corneal limbus region, only 69 cells were identified as limbal epithelial stem cells, corresponding with 0.4% of the total sequenced cells.^43^ Our data presents sequencing information of 5,964 corneal epithelial cells, it is highly possible that there might not have been enough sequencing depth to detect such a rare population. This is an opposing finding to the results presented by Collin et al. 2021, where a major cluster of 893 (4%) limbal progenitor/stem cells was detected in a pool of 21,343 sequenced cells isolated from corneoscleral buttons.^42^

Differential analysis identified the expression of *CAV1*, *HOMER3*, *CXCL14*, and *CPVL* to be exclusive to cell clusters comprising the corneal epithelial stem cell niche (Figure 3). In line with our findings, Collin et al. recently reported the exclusivity of *CXCL14* and *CPVL* to the corneal epithelial stem cell niche.^42^ We further validated the finding of caveolin-1 (*CAV1*) with immunofluorescence, and confirmed it was coexpressed with ΔNp63 in the limbus of corneal tissue and in primary cultured limbal stem cells. These results suggest that caveolin-1 could be used for the identification of epithelial limbal stem cells, with the advantage that it is a cell membrane marker, whereas p63 is expressed in the nucleus. This could open the door to isolating and enriching these cells for regenerative therapies. Furthermore, our study also revealed novel makers specific to transit amplifying cells, namely *CKS2*, *STMN1*, and *UBE2C.* We further validated the finding of cyclin-dependent kinase 2 (*CKS2*) with immunofluorescence, and confirmed it was co-expressed with p63α in the limbus and periphery of human corneas and in primary cultured limbal stem cells, suggesting this is a suitable marker for identifying highly proliferative transit-amplifying cells. Interestingly, primary limbal cells expressing a lower level amount of ΔNp63 and p63α, based on fluorescence intensity, also showed a reduced expression of caveolin-1 and cyclin-dependent kinase 2 respectively (Supplementary Figures S3 and S4).

In line with the study by Collin et al. 2021,^44^ our scRNAseq study shows differential expression of SARS-CoV-2 entry receptors *ACE2*, *TMPRSS2*, *TMPRSS4*^45–47^ in the corneal epithelium, and *NRP1*^48^ in both corneal epithelium and stroma. This results are in line with the hypothesis of the cornea as a potential SARS-CoV-2 entry site (Supplementary Figure S7). Finally, the lack of SARS-CoV-2 entry receptors in the corneal endothelium (Supplementary Figure S7), the most often selectively transplanted corneal layer, supports the safety of donor tissue for endothelial keratoplasty.

It is important to highlight that as previously reported, the corneal epithelium sheds superficial layers when corneas are preserved in media^49^ and it is very likely that this cell atlas does not fully portray the most superficial layer of corneal epithelial cells. Two types of cornea preservation conditions, namely in Optisol-GS medium and organ culture medium have been used for this research, in line with clinical practice in the United States of America and Europe, respectively. No major differences were found between preservation conditions.

Three clusters of the corneal stroma were identified as corneal keratocytes (clusters S0, S1 and S2; Figure 4). Cluster S2 was identified as cells with a key role in extracellular protein secretion to maintain the homeostasis of the stroma. Furthermore, we identified a cluster (S3) of keratocytes transitioning to myofibroblasts, though a fully differentiated myofibroblast phenotype expressing alpha smooth muscle actin was not identified. Interestingly, our analysis did not identify a population of corneal stromal stem cells expressing *ABCB5* and *ABCG2*, as previously identified by Funderburgh and colleagues. ^50–52^ Our hypothesis is that the expression of these genes is induced by the primary expansion of corneal keratocytes and is not found in the cornea.^50–52^ Finally, differential gene expression analysis identified that the expression of *NNMT* was exclusive to the corneal stroma, suggesting this could be used as a novel marker to identify corneal stromal cells.

Two cell clusters were identified forming the corneal endothelium (Figure 5). Both clusters showed high expression of corneal endothelial markers such as CD166 (*ALCAM*), sPrdx1 (*PRDX1*), ZO-1 (*TJP1*), or *SLC4A11*, confirming the endothelial phenotype. Nevertheless, expression of *CD200*, reported as corneal endothelial cell marker in a previous study,^53^ was not detected in our dataset, suggesting it might not be a specific marker for corneal endothelial cells. Despite sharing great similarity, one of the clusters (En1) expressed more extra cellular matrix proteins, suggesting that these cells could play a crucial role on maintaining the Descemet’s membrane. Cell cluster En0 showed lower expression of tight junction proteins and focal adhesions, suggesting these cells could preferentially migrate upon corneal endothelial damage and contribute to tissue repair via migration or cytosolic expansion. Interestingly, our study did not detect a population of precursor endothelial cells, as discussed in previous studies.^54, 55^ Nevertheless, both corneal endothelial cell clusters detected retained expression of *PITX2*, a marker associated with neural crest-derived corneal endothelial cell precursors.^40, 41^ These data suggests that it is highly possible that a progenitor-like state is not exclusive to the peripheral corneal endothelium but to cells across the endothelium.

Interestingly, no corneal endothelial fibroblasts were detected in this study, indicated by the lack of expression of fibroblastic markers α-smooth muscle actin (*ACTA2*) and *CD44* (Supplementary Figure S6), which is in contrast to a previous report.^42^ It is likely that the bulk enzymatic corneoscleral tissue desegregation performed by Collin et al., affected the susceptible corneal endothelial cells, causing endothelial cells transcriptomic bias portrayed in scRNAseq dataset, as showed in other studies.^56, 57^ We performed a more gentle approach to obtain single endothelial cells for sequencing, first dissecting the tissue and mechanically stripping the endothelium from the cornea, and then treating the endothelial cells with a shorter enzymatic digestion.

Overall, this study provides significant information to help understand the heterogeneity of the healthy human cornea, as well as understanding the gene expression stratification across cells present in the same corneal layer, while providing novel markers to identify specific cell types. Moreover, this transcriptomic cell atlas offers a baseline for future studies with the aim of regenerating corneal tissue or further developing corneal cell replacement therapies.

## MATERIALS AND METHODS

### Ethical statement

This study was performed in compliance with the tenets of the Declaration of Helsinki. Ten human donor corneas (Table 1) deemed unsuitable for transplantation were obtained from the Saving Sight Eye Bank (Kansas City, MO, USA) and the ETB-BISLIFE Multi-Tissue Center (Beverwijk, the Netherlands) were used for this study. Both male and female donor corneas, with ages ranging from 22 to 79 years and preserved either in Optisol-GS at 4°C or in Organ culture media at 31°C were used. Organ culture media comprised the following: minimum essential medium supplemented with 20 mM HEPES, 26 mM sodium bicarbonate, 2% (v/v) newborn calf serum (Thermo Fisher Scientific), 10 IU/mL penicillin, 0.1 mg/mL streptomycin and 0.25 μg/mL amphotericin. The tissues used for scRNAseq had no history of ocular disease, chronic systemic disease or infection such as HIV or hepatitis B.

### Manual dissection of the corneal tissue

Cells were isolated from six corneas that were manually dissected to separate the epithelial, stromal and the endothelial layers. The corneas were vacuum fixed in a punch base (e.janach) endothelial-cell side up, stained with trypan blue solution (0.4%) for 30 s and washed with BSS ophthalmic irrigation solution. The corneal endothelium was then gently lifted following the Schwalbe line using a DMEK cleavage hook (e.janach) and fully stripped using angled McPherson tying forceps. The remaining tissue was trephined using a 9.5 mm Ø Barron vacuum punch (Katena) in order to separate the epithelium and stroma from the scleral ring. The remaining two corneas were used for limbus isolation. A surgical scalpel was used to cut the limbus into approximately 1 × 2 mm fragments, which were then rinsed with PBS.

### Tissue dissociation to single cells

The six manually dissected corneal tissues were enzymatically treated to obtain single cell suspensions. The stripped corneal endothelium was incubated with 2 mg/mL collagenase (Sigma) solution in human endothelial serum free media (SFM) (Thermo Fisher Scientific) for 1–2 h at 37°C followed by a 10 min incubation with Accutase (Thermo Fisher Scientific) at 37°C to obtain a single cell suspension. The cells were centrifuged for 5 min at 800 × g and resuspended in 0.5 mL of human endothelial SFM media.

The corneal epithelium–stroma tissues were treated with 1.5 mg/mL collagenase and 0.2 mg/mL bovine testis hyaluronidase (Sigma) in DMEM/F12 (Thermo Fisher Scientific) for approximately 1 h at 37 °C. The released corneal epithelium was suctioned with a P1000 micropipette and further treated with Accutase for 10 min at 37°C to obtain a single cell suspension of corneal epithelial cells. The cells were centrifuged for 5 min at 800 × g and resuspended in 0.5 mL of DMEM/F12. The remaining corneal stroma was treated with 1.5 mg/mL collagenase and 0.2 mg/mL bovine testis hyaluronidase in DMEM/F12 for 5–7 h to obtain a single cell suspension of corneal keratocytes. After the incubation, the cells were centrifuged for 5 min at 800 × g and resuspended in 0.5 mL of DMEM/F12 media.

The limbus fragments isolated from the two corneas for limbal isolation were digested with four trypsinization cycles. In each cycle, the limbus fragments were incubated in 10 mL of 0.05% trypsin/0.01% EDTA (Thermo Fisher Scientific) at 37°C for 30 min. The trypsin containing dissociated cells were transferred to a 50 mL centrifuge tube containing 10 mL DMEM/F12 media and centrifuged for 5 min at 300 × g, after which the cells were resuspended in 0.5 mL of DMEM/F12 media. The undigested limbal fragments were placed again in 10 mL of 0.05% trypsin/0.01% EDTA for the following trypsinization cycle. The cells isolated from the limbus of the two donor corneas were pooled for the following steps.

### Methanol cell fixation

After dissociation, the single cell suspensions were passed through a 100 μm Ø cell strainer, centrifuged for 5 min at 300 × g and resuspended in 1 mL ice-cold PBS to eliminate any medium remnants. Next, the cells were again centrifuged for 5 min at 300 × g and resuspended in ice-cold PBS at a ratio of 200 μL PBS per 1×10^6^ cells followed by the dropwise addition of ice-cold methanol, at a ratio of 800 μL per 1×10^6^ cells. The cells were stored at -80°C until sequencing.

### Single-cell RNA sequencing (scRNAseq)

Single-cell mRNA sequencing was performed at Single Cell Discoveries according to standard 10x Genomics 3’ V3.1 chemistry protocol. Prior to loading the cells on the 10x Chromium controller, cells were rehydrated in rehydration buffer. Cells were then counted to assess cell integrity and concentration. Approximately 9,000 cells from corneas 01, 02, 03 and 04 (3,000 cells per corneal layer), 3,000 cells from corneas 05 and 06 from a 1:1:1 cell layer pool, and 3,000 cells from corneas 07 and 08 (limbal samples) were loaded and the resulting sequencing libraries were prepared following standard 10x Genomics protocol. The DNA libraries were paired end sequenced on an Illumina Novaseq S4, with a 2x150 bp Illumina kit.

### Bioinformatic analysis of scRNA-seq data

BCL files resulting from sequencing were transformed to FASTQ files with 10x Genomics Cell Ranger mkfastq. FASTQ files were mapped with Cell Ranger count. During sequencing, Read 1 was assigned 28 base pairs, and were used for identification of the Illumina library barcode, cell barcode and UMI. R2 was used to map the human reference genome GRCh38. Filtering of empty barcodes was done in Cell Ranger. The data from all samples were loaded in R (version 3.6.2) and processed using the Seurat package (version 3.2.0).^58^ More specifically, cells with at least 1000 UMIs per cell and less than 20% mitochondrial gene content were retained for analysis. The data of all 10x libraries was merged and processed together. The merged dataset was normalized for sequencing depth per cell and log-transformed using a scaling factor of 10,000. The patient effect was corrected using the integration function of Seurat and used for dimensionality reduction and clustering of all cells or cells selected per layer. Cells were clustered using graph-based clustering and the original Louvain algorithm was utilized for modularity optimization. The differentially expressed genes per cluster were calculated using the Wilcoxon rank sum test and used to identify cell types. Putative doublets were computationally identified using scDblFinder (v1.2.0).^59^

### Primary culture of limbal cells

Human primary limbal cells were harvested from corneal tissue of cadaveric donors (ages ranging from 36 to 79 years) with informed consent. Human limbal cells were cultured as previously described.^60, 61^ In short, a surgical scalpel was used to cut the limbus of the corneas into approximately 1 × 2 mm fragments, which were then rinsed with PBS. The limbus fragments were then incubated in 10 mL of 0.05% trypsin/0.01% EDTA (Thermo Fisher Scientific) at 37°C for 30 min. The trypsin containing dissociated cells were transferred to a 50 mL centrifuge tube containing 10 mL culture media and centrifuged for 5 min at 300 × g, after which the cells were resuspended in 0.5 mL of culture media. The undigested limbal fragments were placed again in 10 mL of 0.05% trypsin-EDTA for another trypsinization cycle. Culture medium consisted of 2:1 mixture of DMEM/F12 media (Thermo Fisher Scientific) supplemented with 2 mM GlutaMAX (Thermo Fisher Scientific), 10% fetal bovine serum (Thermo Fisher Scientific), 125 IU/L insulin (Humulin R, Lilly), 0.2 mM adenine (Merck), 1,1 μM hydrocortisone (Merck), 8.5 ng/ml cholera toxin (Sigma), 2 nM triiodothyronine (Sigma), 10 ng/ml epidermal growth factor (Amsbio), and 100 IU/mL penicillin-streptomycin (Thermo Fisher Scientific). The limbal cells were plated on a feeder layer of lethally irradiated 3T3-J2 fibroblasts (fibroblast feeder-layer density 40,000 cells/cm^2^). The 3T3-J2 fibroblast immortalized cells line was a kind gift of Prof. Howard Green (Harvard Medical School, Boston, MA, USA). When confluent, limbal epithelial stem cells were passaged by 0.05% trypsin/0.01% EDTA (Thermo Fisher Scientific) treatment and seeded at a density of 15,000 cells/cm^2^ in a Nunc chamber slide (Thermo Fisher Scientific).

### Immunofluorescence

Two human donor corneas and primary cultured limbal cells were used for immunofluorescence analysis. The corneas were cut in half transversally, embedded in a cryomold containing Tissue-Tek O.C.T. compound, snap frozen in liquid nitrogen, and stored at -80°C until sectioning. For sectioning, 10 μm consecutive sections were cut on an adhesive cryofilm type 3C(16UF) using a modified Kawamoto method^62^, to help preserve the morphology of the tissue during sectioning. The sections were left to dry for 10 min prior to use. The corneal sections and the primary limbal cells cultured on a chamber slide were fixed with 4% paraformaldehyde, permeabilized with 0.1% Triton X-100 in PBS for 10 min and blocked with 2% BSA solution in PBS for 1 h followed by overnight incubation at 4**°**C with primary antibodies diluted in 2% BSA blocking solution: mouse monoclonal [1F7G5] anti-CKS2 (1:100 dilution; Thermo Fisher Scientific; 37-0300), rabbit polyclonal anti-P63 (p63α) (1:100 dilution; Cell Signaling Technology; 4892), rabbit polyclonal anti-caveolin1 (1:300 dilution; Abcam; ab2910), and mouse monoclonal [4A4] anti-P63 (ΔNp63) (1:100 dilution; Abcam; ab735). The tissue sections and primary limbal cells were washed five times and incubated with secondary antibodies diluted in 2% BSA blocking solution, goat anti-rabbit A488 (1:300 dilution; Thermo Fisher Scientific) or goat anti-mouse A568 (1:300 dilution; Thermo Fisher Scientific), for 50 min at ambient temperature in the dark. Cell nuclei were stained with 0.5 μg/mL DAPI for 10 min. The samples were washed five times in PBS, mounted with coverslips with Fluoromount G mounting medium (Thermo Fisher Scientific) and examined on a Nikon Eclipse Ti-E inverted microscope equipped with a X-Light V2-TP spinning disk (Crest Optics).

## Funding

This research was funded by Chemelot InSciTe under the EyeSciTe consortium.

## Author contributions

P.C.: experimental design, tissue acquisition, tissue dissection and dissociation to single cells, microscopy imaging for immunofluorescence experiments, experimental data analysis, manuscript and figures preparation. N.G.: bioinformatics analysis, data collection, manuscript and figures preparation. J.A.D.: experimental data analysis. E.S.: experimental design, tissue acquisition, tissue dissection and dissociation to single cells, experimental data analysis. A.J.H.v.V.: limbal tissue dissection and dissociation to single cells, immunofluorescence experiments, manuscript preparation. V.L.S.L., M.M.D., and R.M.M.A.N.: fund acquisition, experimental design, tissue acquisition, manuscript and figures preparation, and supervised the work. All authors ensured that questions on the accuracy or integrity of all parts of the study were appropriately researched and resolved.

## Competing interests

The authors declare no conflict of interest.

## Acknowledgements

The authors thank Annika Jeschke and Timo Rademakers (Maastricht University) for the assistance in the immunofluorescence experiments and the sample cryosectioning. The authors thank Single Cell Discoveries (Utrecht, the Netherlands) for the single cell sequencing services provided.

The data regarding this study will be deposited to the Gene Expression Omnibus.

---

**Supplementary Figure S1.**
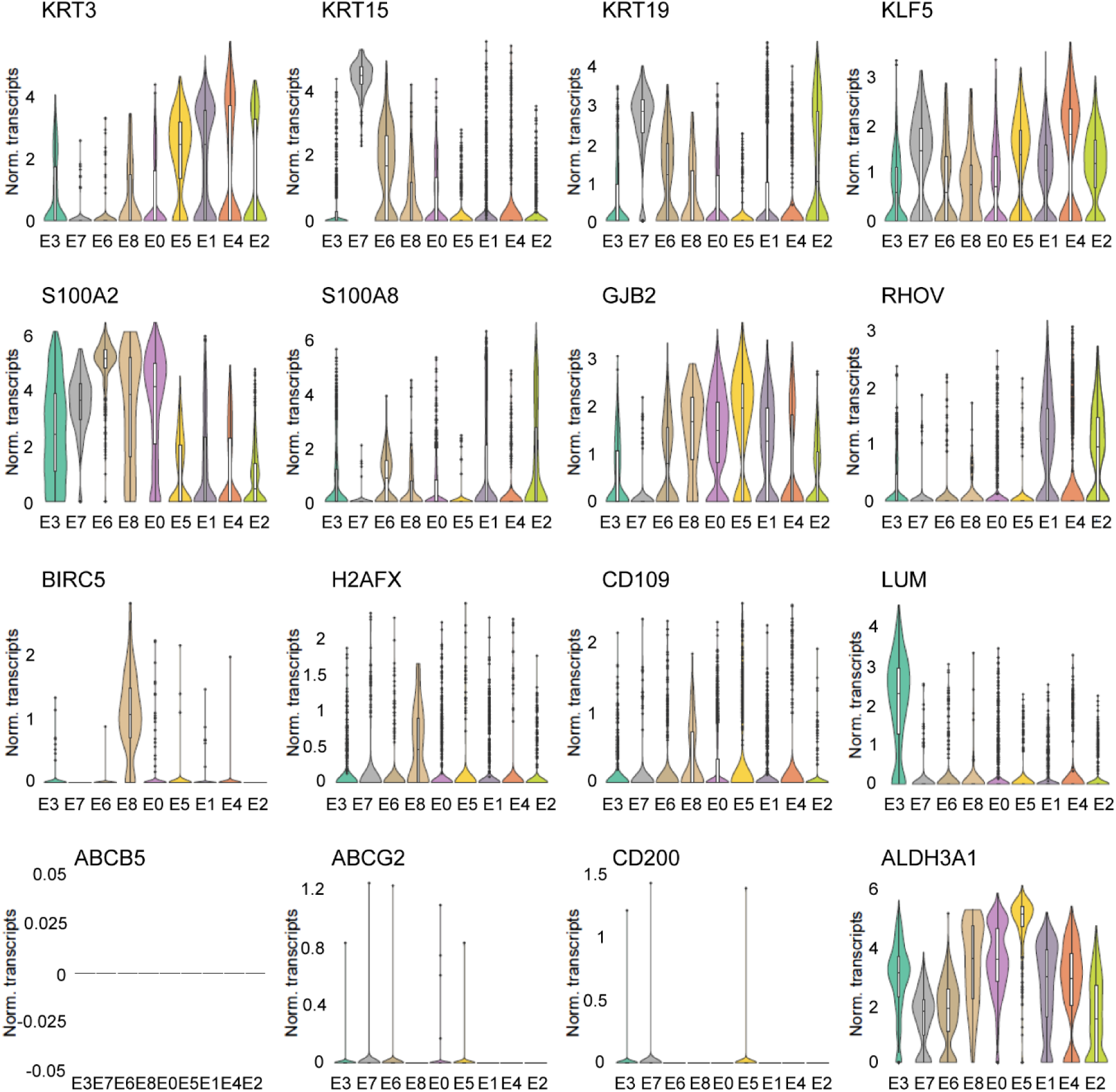
Violin plots show additional marker genes for the identification of corneal epithelial clusters.

**Supplementary Figure S2.**
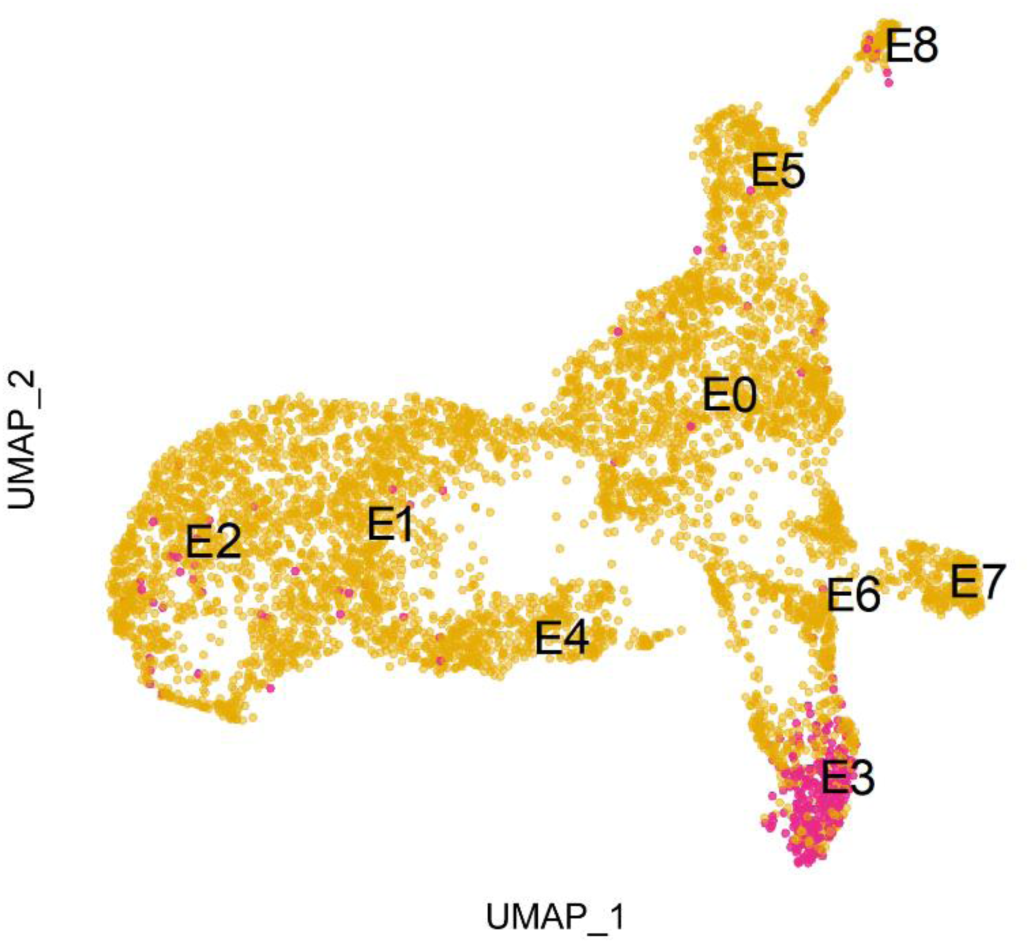
Putative doublets were computationally identified using scDblFinder (v1.2.0) and epithelial cell cluster E3 was identified as a cluster of stromal-epithelial doublets.

**Supplementary Figure S3.**
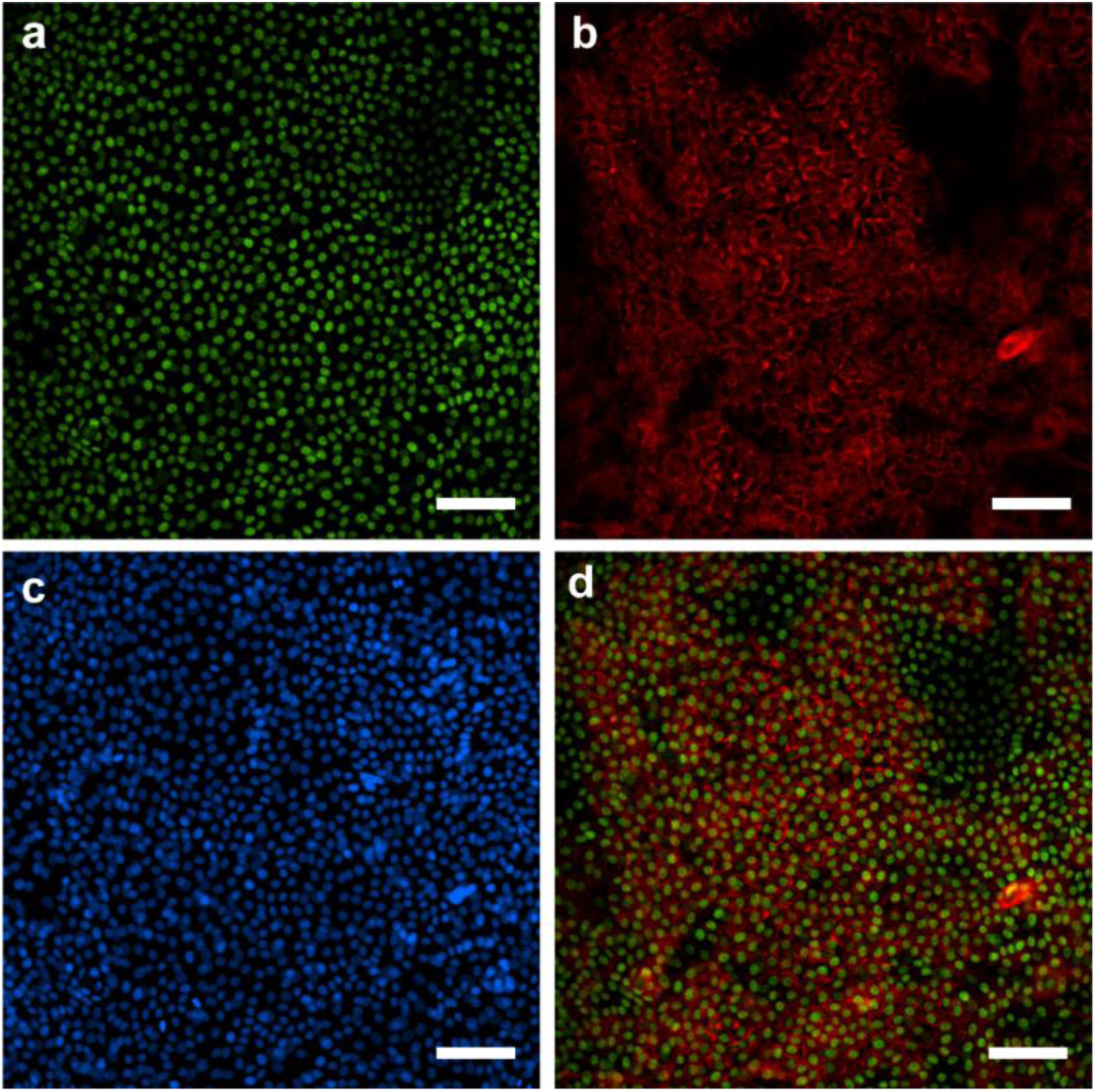
Immunofluorescence analysis of caveolin-1 (*CAV1*) expression in primary cultured human corneal limbal epithelial stem cells. Limbal epithelial stem cells expressing ΔNp63 (a, green) also showed expression of caveolin-1 (b, red), as observed in the image overlay (d). Cell nuclei were stained with DAPI (c). These results suggest caveolin-1 could be a selective marker for corneal limbal stem cells. Scale bar is 100 μm.

**Supplementary Figure S4.**
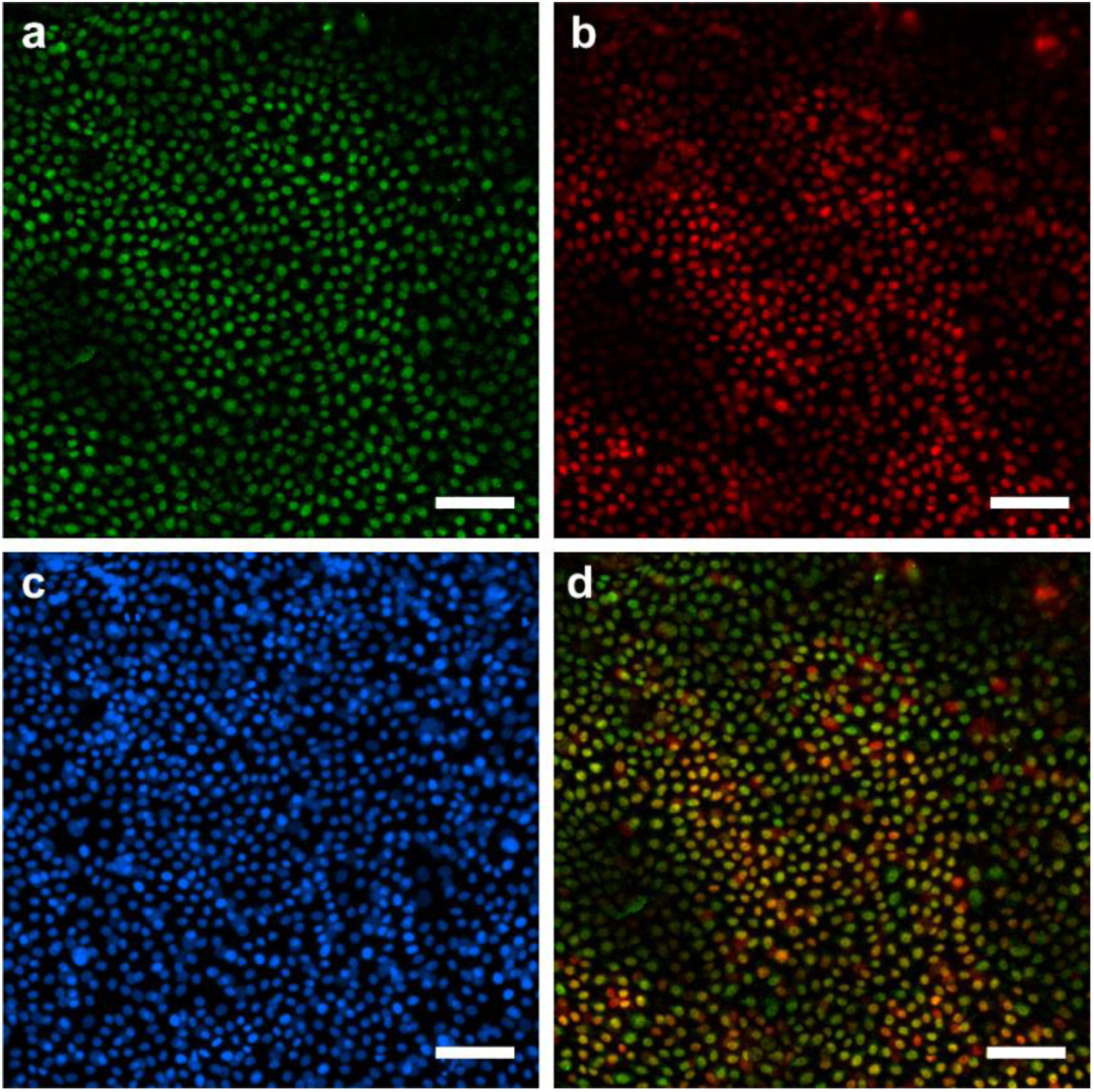
Immunofluorescence analysis of cyclin-dependent kinase 2 (*CKS2*) expression in primary cultured human corneal limbal epithelial stem cells. Limbal epithelial stem cells expressing p63α (a, green) also showed expression of *CKS2* (b, red), as observed in the image overlay (d). Cell nuclei were stained with DAPI (c). Scale bar is 100 μm.

**Supplementary Figure S5.**
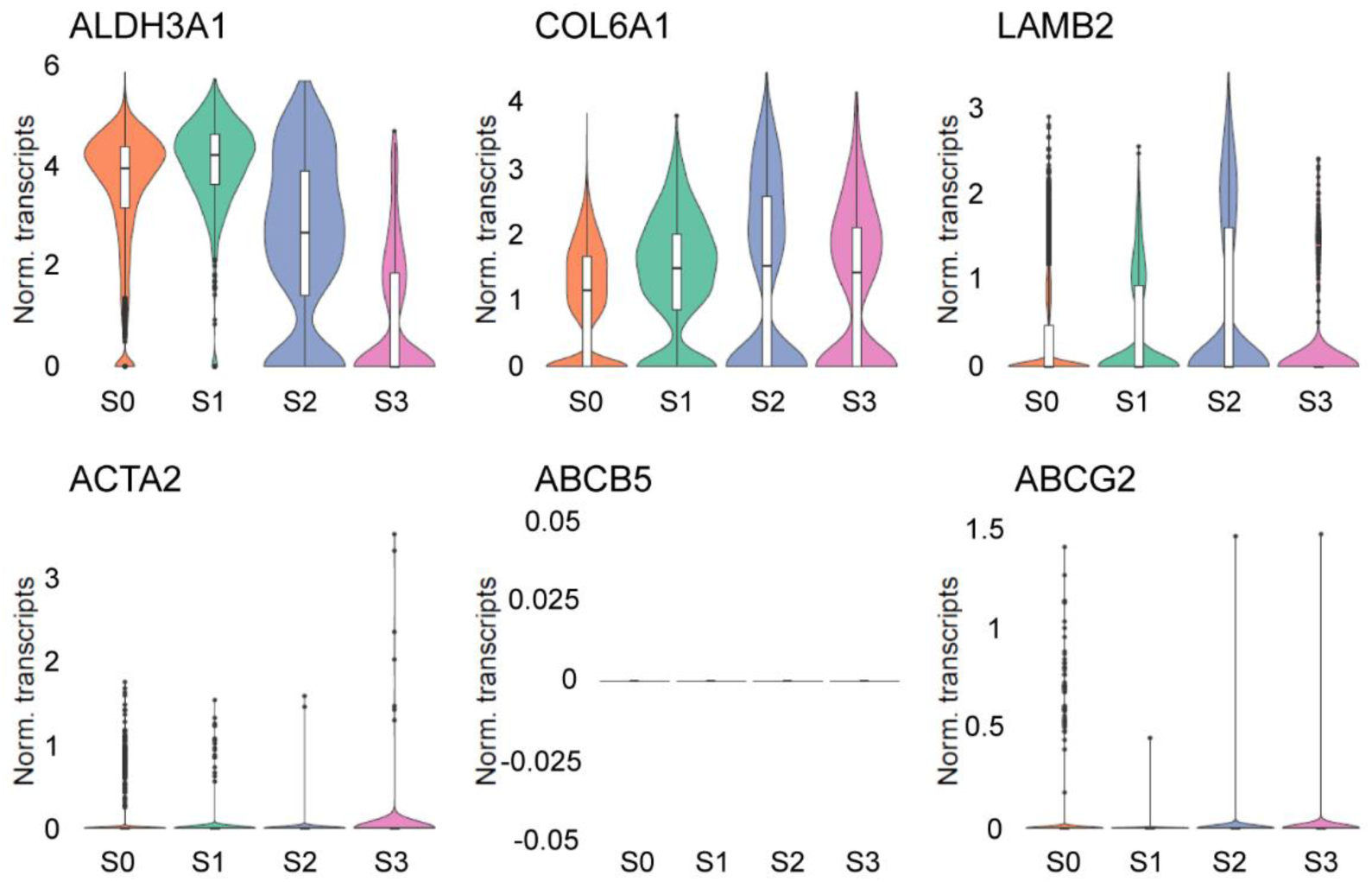
Violin plots show additional marker genes for the identification of corneal stromal clusters.

**Supplementary Figure S6.**
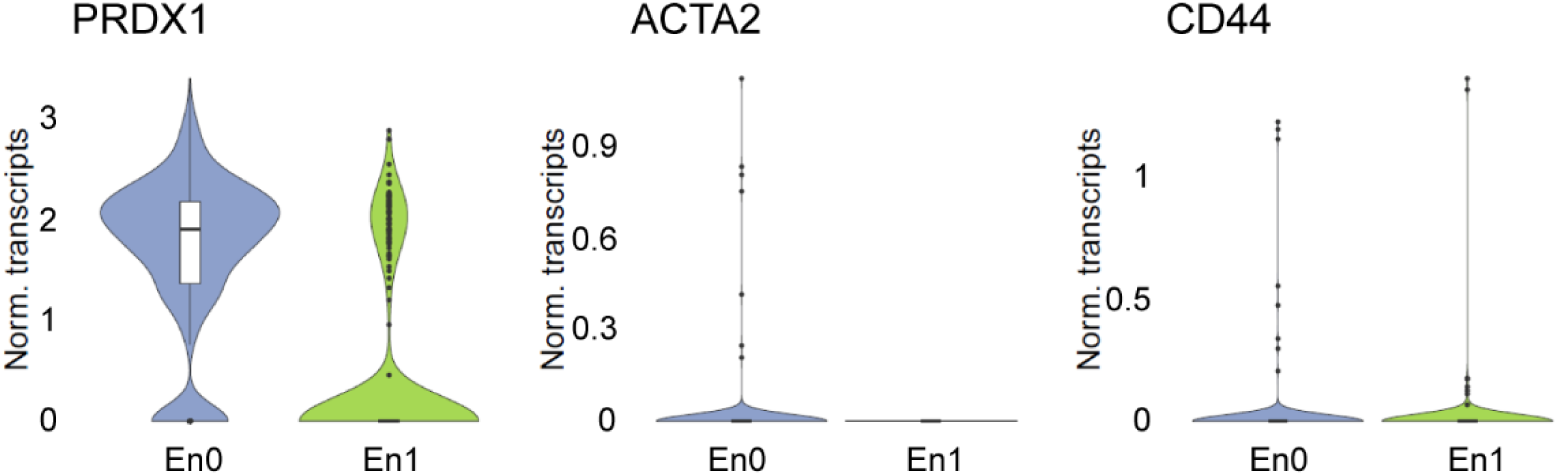
Violin plots show additional marker genes for the identification of corneal endothelial clusters.

**Supplementary Figure S7.**
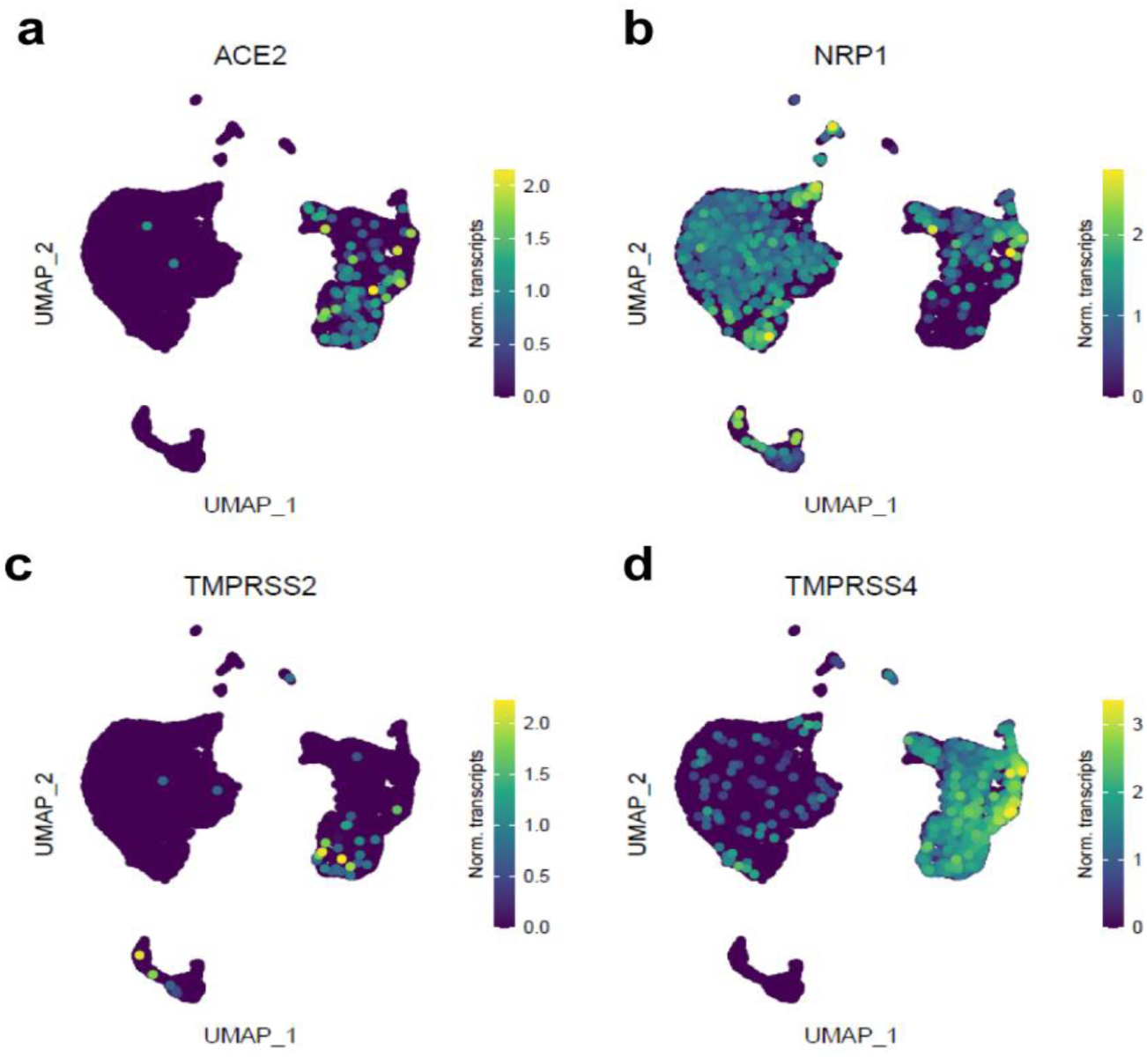
Single cell transcriptome expression level UMAP of SARS-CoV-2 cell receptor *ACE2* (a) and receptor-associated *NRP1* (b), *TMPRSS2* (c), and *TMPRSS4* (d) protein expressing genes.

## Notes

### Competing Interest Statement

The authors have declared no competing interest.

